# TL1A-activated T cells remodel the rectal mucosa in Crohn’s disease patients with perianal fistulizing disease

**DOI:** 10.1101/2025.06.26.657455

**Authors:** Victoria Gudiño, Jae-Won Cho, Berta Caballol, Ana M. Corraliza, Marisol Veny, Isabella Dotti, Livia Moreira Genaro, Ángela Sanzo-Machuca, Elisa Melón-Ardanaz, M. Carme Masamunt, Miriam Esteller, Iris Teubel, Lisseth Robbins, Ángel Giner, Cristina Prieto, Elena Ferrer, Raquel Franco-Leal, Albert Martin-Cardona, Carme Loras, Maria Esteve, Jordi Rimola, Agnès Fernández-Clotet, Ingrid Ordás, Elena Ricart, Julian Panés, Martin Hemberg, Azucena Salas

**Affiliations:** Institut d’Investigacions Biomèdiques August Pi i Sunyer (IDIBAPS), Inflammatory Bowel Disease Unit, Department of Gastroenterology, Hospital Clínic de Barcelona, Barcelona, Spain; Centro de Investigación Biomédica en Red de Enfermedades Hepáticas y Digestivas, CIBEREHD), Madrid, Spain; Gene Lay Institute of Immunology and Inflammation, Brigham and Women’s Hospital, Massachusetts General Hospital and Harvard Medical School, Boston, MA, USA; Inflammatory Bowel Disease Research Laboratory, Gastrocenter, Colorectal Surgery Unit, Department of Surgery, School of Medical Sciences, University of Campinas (UNICAMP), Campinas, São Paulo, Brazil; Facultat de Medicina i Ciències de la Salut, Universitat de Barcelona, Spain; Department of Gastroenterology, Hospital Universitari Mútua Terrassa, Universitat de Barcelona, Terrassa, Spain; BCLC Group, Radiology Department, CDI, Hospital Clinic of Barcelona, University of Barcelona, Barcelona, Spain

**Keywords:** Fistula, single-cell RNA sequencing, Lymphotoxin beta, Fibroblasts

## Abstract

**Background:** Perianal fistulizing disease (PFD) is a complication that affects about 20% of Crohn’s disease (CD) patients whose etiology remains unknown.

**Objectives:** To identify predisposing events driving fistula formation.

**Design:** Rectal biopsies from CD patients with or without PFD (CD+PFD and CD, respectively; n=31) were collected and subjected to single-cell RNA sequencing (scRNA-seq). Functional analyses were conducted using peripheral CD3+ T cells, intestinal tissue explants, primary fibroblasts and 2D-epithelial monolayer cell cultures.

**Results:** The rectal mucosa of CD+PFD patients is imprinted with cellular and transcriptomic alterations specific to PFD and independent of luminal inflammation, potentially driven by TL1A activation in CD4+ T cells. We identified lymphotoxin beta (*LTB* or its functional heterotrimer LTα_1_β_2_) as a novel mediator downstream of TL1A that, along with IL-22, induces a PFD-associated signature in rectal fibroblast and epithelial cells, respectively. This signature includes an increased abundance of fibroblasts, an induction of matrix-degrading enzymes, transcriptomic rewiring of the lamina propria S1 fibroblasts, and an anti-bacterial and immune responses in epithelial cells. Notably, the induction of LTα_1_β_2_ and IL-22 occurs independently of TNF signaling, revealing a new TL1A-LTα_1_β_2_/IL-22 axis that remains active under anti-TNF therapy.

**Conclusion:** Our findings revealed unique cellular alterations in the rectum of CD patients with PFD, highlighting the previously unrecognized involvement of TL1A in mediating this signature and supporting the need for exploring the role of TL1A inhibition as a therapeutic approach for PFD.

## INTRODUCTION

Crohn’s disease (CD) is a chronic remitting and relapsing inflammatory condition of the gastrointestinal tract often complicated by fibro-stenotic, fistulizing, and/or perianal disease. Perianal lesions are present in approximately 12% of patients at diagnosis(1), with their prevalence rising to nearly 20% over the course of the disease(2). Several factors, such as rectal luminal disease, have been associated with an increased risk of developing perianal fistulizing disease (PFD)(3). However, a large proportion of patients with PFD present no rectal involvement, suggesting that the mechanisms driving fistula formation may differ from those supporting luminal inflammation.

The current model used to explain the pathogenesis of PFD involves the transformation of epithelial cells through epithelial-to-mesenchymal transition (EMT) as a response to cytokines such as transforming growth factor beta (TGFβ), tumor necrosis factor (TNF), or interleukin (IL)-13(4–6). This is accompanied by the remodeling of the extracellular matrix (ECM), which facilitates the migration of transformed epithelial cells to the site of fistula formation, as seen by the enhanced expression of matrix-degrading enzymes or metalloproteinases (MMPs) in the fistula compared to intestinal or peri-fistula tissues(7,8). Importantly, experiments using cells from the fistula tract revealed an expansion of CD4+ over CD8+ T cells(6), as well as a heightened fibroblast response to IL-13 or TGFβ compared to fibroblasts from non-fistulizing tissue(4). However, these observations are largely based on analyses from fistula-derived tissues, which are more likely to reflect the end-stage of the disease rather than the early mechanisms driving fistula initiation or progression.

To address this limitation, we analyzed the rectal mucosa of CD patients with (CD+PFD) and without (CD) perianal disease, generating a comprehensive single-cell atlas of the rectal cellular populations and states. Considering the increased risk of fistula development in patients with rectal involvement and the possibility that fistulae may originate from both the anal canal and the rectum, we hypothesized that alterations in the rectum may precede and contribute to the development of PFD, through mechanisms potentially independent from those driving luminal inflammation.

Our data reveals that the transcriptomic rewiring of the rectal mucosa can occur independently of luminal inflammation and is, at least in part, mediated by the activation of CD4+ T cells in response to tumor necrosis factor-like ligand 1A (TL1A). Herein, we provide novel insights into the pathophysiology of perianal disease, defining a distinct stromal and epithelial signature associated with PFD and identifying TL1A along with lymphotoxin β and IL-22 as key mediators of these alterations with potential therapeutic value.

## METHODS

Detailed methods are outlined in the online supplementary information.

## RESULTS

### Defining a PFD signature in the rectal mucosa

To investigate whether changes in the rectum could contribute to the development of fistulae, we analyzed a cohort of CD patients with (CD+PFD; n=19) or without (CD; n=12) PFD (Figure 1A, Supplementary Table 1). Biopsies were freshly processed for scRNA-seq as described(9). We obtained the transcriptomes from 81,370 high quality cells, which we annotated into 5 subsets: epithelium, stroma, T cells, B and plasma cells, and myeloid cells (Figure 1B). These were then subclustered to identify cell types at a higher resolution (Supplementary Figure 1A, Supplementary Table 2). A comparison of the cell abundances between the two groups revealed a significant increase in the number of lamina propria S1 (*ABCA8)*, bottom-crypt S3 (*OGN*) and immediate-early response (IERs) fibroblasts (*FOS*) and glia (*S100B*) in CD+PFD compared to CD patients (Figure 1C). In contrast, CD samples showed an increase in the abundance of pericytes (*RSG5*), IgG plasma cells, plasmablasts (*DERL3*) and neutrophils (*PROK2*). Importantly, changes in all the latter populations could be attributed to differences in the severity of inflammation within samples from both groups, rather than changes associated with the presence of PFD. Indeed, each group contained samples from endoscopically inflamed rectum (6/19 in the CD+PFD group -CD+PFDinf- and 5/12 in the CD group -CDinf-; Figure 1D, Supplementary Figure 1B, C). In addition, some of the patients in the CD+PFD group without rectal inflammation at the time of sampling had previous rectal involvement (CD+PFD with rectal mucosa in remission, CD+PFDre, n=6). Given the impact that current and even previous disease has on the transcriptome of the intestinal mucosa(10), we focused our analysis on samples from uninvolved rectum (patients with no current or previous rectal inflammation) from CD+PFD (CD+PFDun, n=7) and CD (CDun, n=7) patients. Of note, none of the patients in the CDun group developed PFD even after 5 years since inclusion, confirming their stable non-fistulizing phenotype. Interestingly, when comparing CD+PFDun and CDun groups, we confirmed the increase in different fibroblast subsets and glia in the fistulizing group, along with myeloid, IgG plasmablasts and some T cell types (Figure 1E), suggesting that not all of them are driven by inflammation. Beyond changes in cell abundances, differentially expressed gene (DEG) analysis between these groups revealed significant changes in gene expression across lymphoid, epithelial, and stromal cells (Figure 1F, Supplementary Figure 1D). In sum, these results indicate that the rectal mucosa of PFD patients, even in the absence of ongoing or previous inflammation, is imprinted with a unique transcriptional profile that involves lymphocytes, various epithelial cell types, and fibroblasts.

**Figure 1:**
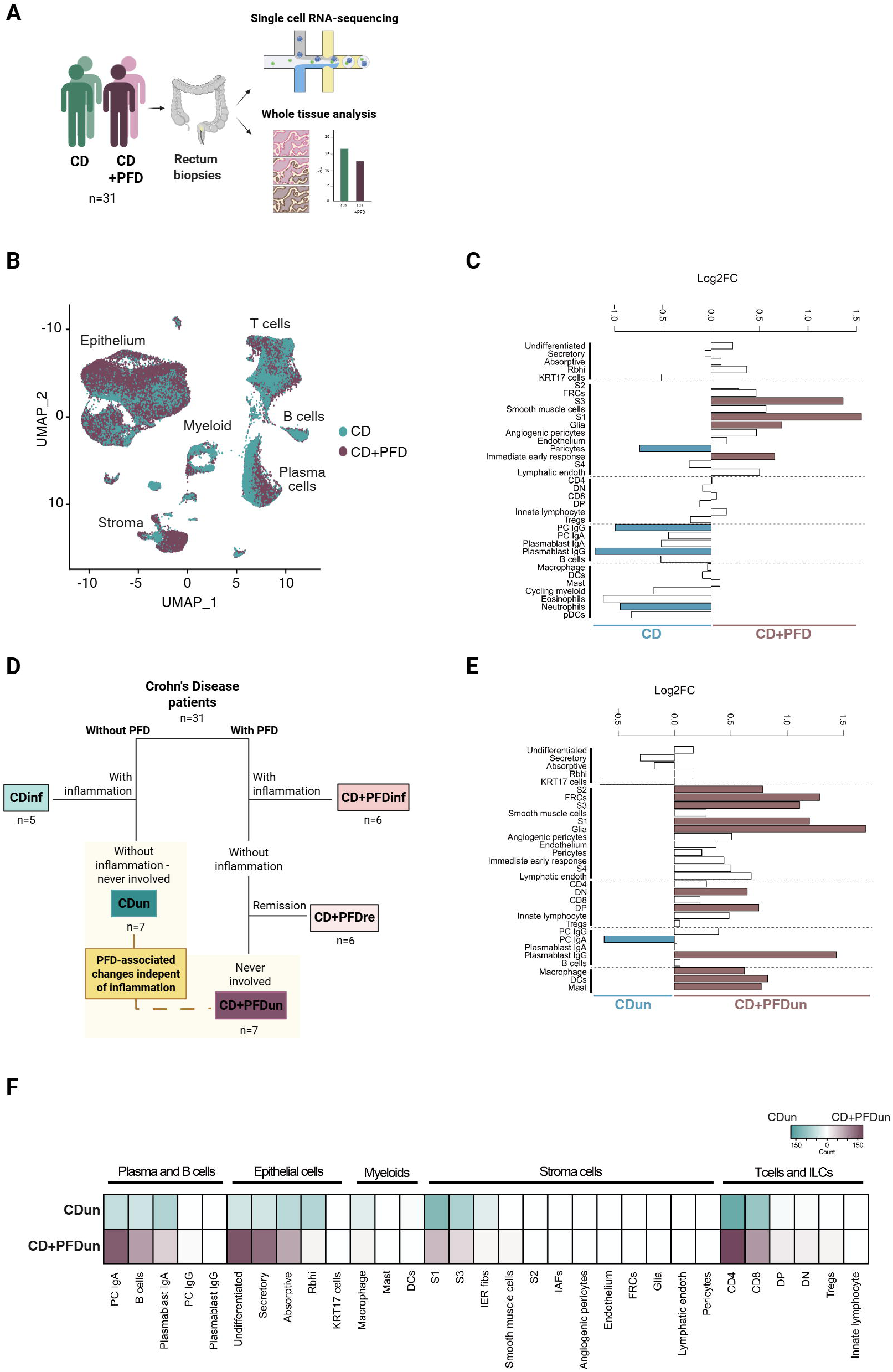
Patients with PFD present cellular and transcriptomic alterations in the rectal mucosa. **A:** Schematic representation of the study. **B:** Uniform Manifold Approximation and Projection (UMAP) representation of the scRNA-seq dataset colored by patient group. **C:** Cell abundance analysis comparing CD and CD+PFD patients. **D:** Chart indicating the categorization of patients from the cohort based on the presence of PFD and luminal inflammation of the rectum. **E:** Cell abundance analysis comparing CDun and CD+PFDun patients. For **C** and **E**, cell frequency differences are represented as the logarithm of fold change in base 2 (Log2FC), and colored bars indicate significant differences (|FC| ≥ 1.5 and adjusted p-value < 0.05). Fisher’s exact test, Benjamini-Hochberg correction. **F:** Heatmap representing the number of differentially expressed genes (DEGs) between CDun and CD+PFDun patients. Only cell types with cell counts above 10 are shown.

### Changes in lamina propria fibroblasts from PFD patients occur independently of TGFβ

Cell abundance analysis pointed to fibroblasts as significantly altered in PFD patients regardless of their inflammatory state. Lamina propria S1 fibroblasts showed the highest number of DEGs when comparing CDun with CD+PFDun patients (Figure 1F, Figure 2A, Supplementary Table 3). Pathway analysis suggests a metabolic reprogramming of CD+PFDun S1 fibroblasts, with pathways related to oxidative phosphorylation and lipid metabolism being significantly enriched (Supplementary Figure 2A). Intriguingly, TGFβ signaling, which has previously been suggested to play a role in the pathophysiology of PFD(4,5), was significantly downregulated in S1 fibroblasts from the CD+PFDun group (Figure 2B). To confirm this, we stimulated primary intestinal fibroblasts with TGFβ and measured the expression of several genes we found upregulated in fibroblasts from CD+PFDun patients. Indeed, TGFβ significantly downregulated the expression of some of those genes, while it induced the expression of *CTGF,* the second-most downregulated gene in the CD+PFDun group (Figure 2C). Interestingly, TGFβ also decreased the expression of *MMP1*, *MMP3* and *CHI3L1*, which previous studies found upregulated in perianal fistula tracts(7,8,11). In agreement with the scRNA-seq data, bulk RNA from the rectal biopsies showed not only the significant upregulation of *TFPI2,* though not *EMILIN1* and *PLAC9*, but also the downregulation of *CTGF* in CD+PFDun biopsies (Figure 2D, Supplementary Figure 2B). Interestingly, we also observed a significant upregulation of *MMP1*, *MMP3* and *CHI3L1* in bulk RNA from CD+PFDun samples, validating their association with PFD (Figure 2D).

**Figure 2:**
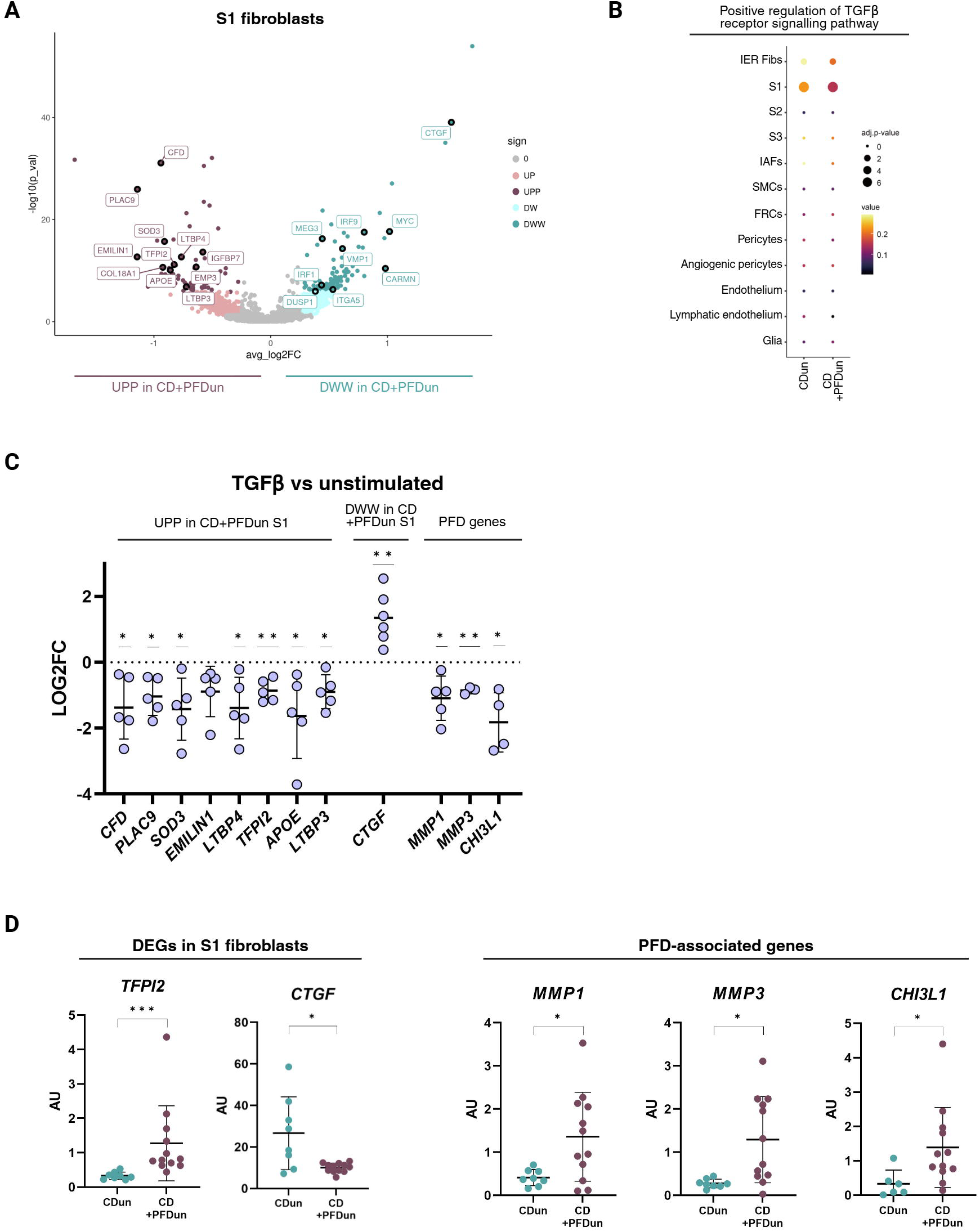
Transforming growth factor beta (TGFβ) signaling is downregulated in the S1 fibroblasts of CD+PFDun patients. **A:** Volcano plot representing the differentially expressed genes (DEGs) in the lamina propria S1 fibroblasts between CDun and CD+PFDun patients. A two-sided Wilcoxon rank sum test was applied. Genes with an adjusted p-value < 0.05, and an avgLog2FC ˃ log2 (1.2) or avgLog2FC ˂ -log2 (1.2) were considered up-(UPP) or down-regulated (DWW), respectively. Genes with a nominal p-value ˂ 0.05 are also indicated in light colors if avgLog2FC ˃ log2 (1.2) (UP) or avgLog2FC ˂ -log2 (1.2) (DW). **B:** Dot plot representing the activity of the “positive regulation of TGFβ receptor signaling pathway” process in each mesenchymal cell type comparing CDun and CD+PFDun patients. Dot size indicates the adjusted p-value, while color scale indicates the arbitrary value of the pathway generated by the AddModuleScore function in the Seurat package. The Wilcoxon-rank sum test (two-sided) with Holm correction was applied to compare CDun and CD+PFDun patients. **C:** qRT-PCR of primary intestinal fibroblasts stimulated with TGFβ (5ng/ml) for 24 hours. Data is shown as a logarithm of the fold change in base 2 (Log2FC) comparing TGFβ-stimulated *vs* unstimulated fibroblasts. One sample t-Test, parametric, * = p < 0.05, ** p = < 0.01, n = 5. **D:** Whole-tissue qRT-PCR analysis of rectal biopsies from CDun (n = 8) and CD+PFDun (n = 12) patients; Mann-Whitney test, * = q ≤ 0.05, ** = q ≤ 0.01, **** = q ≤ 0.0001. AU = arbitrary units.

Finally, we explored whether this signature was also observed in the context of mucosal inflammation. While there were no DEGs between CDinf and CD+PFDinf patients in the S1 fibroblasts (Supplementary Figure 2C), a fibroblast-specific TGFβ response signature was also downregulated in CD+PFDinf compared to CDinf, suggesting that this might be a specific PFD trait (Supplementary Figure 2D). In summary, we describe a transcriptional signature expressed by fibroblasts and associated with the PFD phenotype. This signature was not driven by TGFβ signaling, suggesting alternative mechanisms may be at work.

### CD4+ T cell-expressed lymphotoxin β imprints a PFD signature on intestinal fibroblasts

We next hypothesized that immune-derived signals, independent of TGFβ, may be acting on the stromal compartment of PFD patients. Within immune cells, T cells, particularly CD4+ T cells, were the cell type with the highest number of DEGs when comparing CDun and CD+PFDun patients (Figure 1F, Supplementary Table 3). Among DEGs, *LTB*, encoding for the lymphotoxin β chain, presented the highest fold change increase across all T cell subsets in patients with perianal disease (Figure 3A and Supplementary Figure 3A). This lymphotoxin subunit forms a membrane-bound heterotrimer with LTα (LTα_1_β_2_) and signals through the LTβ receptor (*LTBR*)(12). *LTBR* is expressed by stromal and epithelial cells, in addition to some myeloid subsets (Figure 3B). We then investigated *LTB-LTBR* interactions across all cell types using CellPhoneDB(13). In agreement with the expression pattern of *LTBR*, cells expressing *LTB* interacted with epithelial, mesenchymal and myeloid cells, being the strength and interaction of *LTB-LTBR* higher in CD+PFDun compared to CDun patients (Figure 3C). Based on this, we hypothesized that *LTB* in its heterotrimer form LTα_1_β_2_ could be driving the transcriptional signature we observed in fibroblasts from CD+PFDun patients.

**Figure 3:**
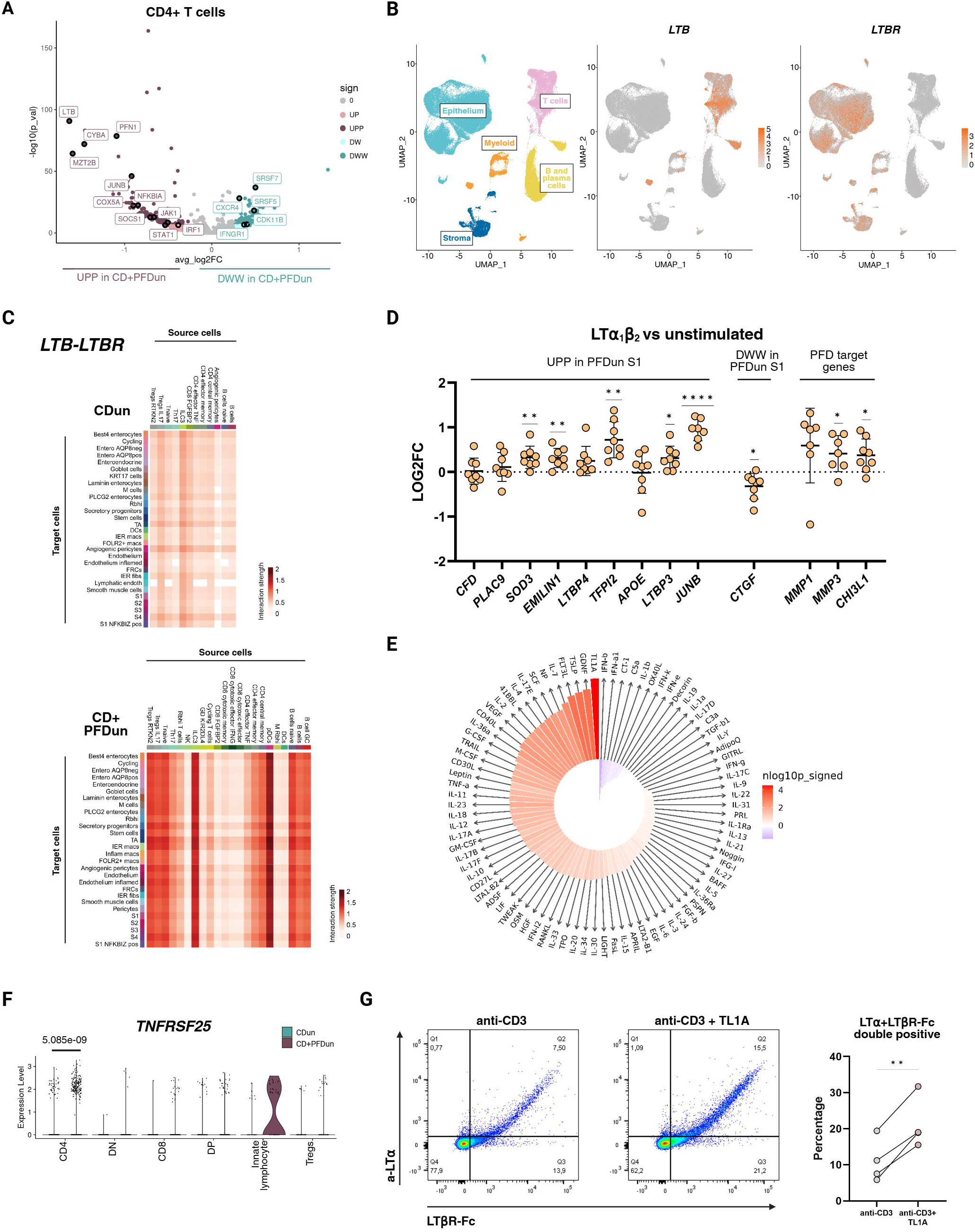
LTα_1_β_2_ derived from TL1A-activated T cells induces the PFD-associated signature in the stroma. **A:** Volcano plot representing the differentially expressed genes (DEGs) in the CD4+ T cells between CDun and CD+PFDun patients. A two-sided Wilcoxon rank sum test was applied. Genes with an adjusted p-value < 0.05, and an avgLog2FC ˃ log2 (1.2) or avgLog2FC ˂ -log2 (1.2) were considered up-(UPP) or down-regulated (DWW), respectively. Genes with a nominal p-value ˂ 0.05 are also indicated in light colors if avgLog2FC ˃ log2 (1.2) (UP) or avgLog2FC ˂ -log2 (1.2) (DW). **B:** Uniform Manifold Approximation and Projection (UMAP) plot showing the expression of *LTB* and *LTBR* in the whole object. **C:** Heatmaps representing the interactions predicted for *LTB* (source cells) and *LTBR* (target cells) for CDun and CD+PFDun by CellphoneDB analysis. Only significant interactions are represented, and color scale indicated the predicted interaction strength. **D:** qRT-PCR of primary intestinal fibroblasts stimulated with LTα_1_β_2_ (100ng/ml) for 24 hours. Data is shown as a logarithm of the fold change in base 2 (Log2FC) comparing LTα_1_β_2_-stimulated *vs* unstimulated fibroblasts. One sample t-Test, parametric, * = p < 0.05, ** = p < 0.01, n = 7. **E:** Cytokine enrichment plot based on the immune response enrichment analysis(14) conducted with the top fifty genes overexpressed in the CD4+ T cells of CD+PFDun patients. Bar length represents the enrichment score, and the color scale indicates the FDR-adjusted p-value (two-sided Wilcoxon rank-sum test). **F:** Violin plot of *TNFRSF25* expression within the T-cell compartment. The adjusted p-value was generated by DEG analysis. **G:** Flow cytometry plots of peripheral T cells stimulated with anti-CD3 or anti-CD3+TL1A for 48 hours and stained with a-LTα and LTβR-FC. Double-positive cells correspond to LTα_1_β_2_+ T cells. Plots are from a representative replicate, and percentages of LTα_1_β_2_+ T cells comparing the two conditions are shown on the right. Paired t-Test, parametric, ** = p < 0.01, n = 4.

Indeed, primary intestinal fibroblasts stimulated with LTα_1_β_2_ significantly upregulated the expression of *TFPI2*, *EMILIN1*, *SOD3, LTBP3*, *JUNB*, *MMP3* and *CHI3L1* while downregulating *CTGF*, partially reproducing the signature we observed in S1 fibroblasts from CD+PFDun patients (Figure 3D). Modulation of *TFPI2* by LTα_1_β_2_ was also validated at the protein level (Supplementary Figure 3B). Altogether, these results suggest that increased LTα_1_β_2_ expression by CD4+ T cells could be driving the fibroblast signature we observed in CD+PFDun rectal samples.

### A TL1A-LTα_1_β_2_ axis is enhanced in CD+PFDun mucosal T cells

To identify the potential upstream regulators of the CD4+ T cell transcriptional signature in PFD, including the over-expression of *LTB*, we submitted the 50 most significantly regulated genes to the immune dictionary (www.immune-dictionary.org)(14) (Supplementary Table 2). This tool uses single-cell transcriptomic profiles of response to cytokines in murine lymph nodes to perform an immune response enrichment analysis, which infers those potential cytokines that induce the input gene set from a given cell type. This analysis identified TNF-like ligand 1A (TL1A) as the most likely regulator of the CD4+ T cell signature in CD+PFDun samples (Figure 3E). Furthermore, *TNFRSF25*, encoding for the TL1A receptor (DR3), was upregulated in CD+PFDun rectal CD4+ T cells (Figure 3F), further supporting a role for TL1A in inducing the changes observed in the CD4+ T cells of these patients. Conversely, in the context of rectal inflammation (CDun vs CDinf), both *LTB* and *TNFRSF25* were significantly upregulated in CD4+ T cells (Supplementary Figure 3C), with no differences observed in the active mucosa of patients with PFD (CDinf vs CD+PFDinf, Supplementary Figure 3D).

TL1A (*TNFSF15*), which is primarily produced by epithelial cells, was not differentially expressed between groups (Supplementary Figure 3E). This cytokine is known for its role in T cell activation, particularly Th1(15). However, to our knowledge, a link between TL1A and LTα_1_β_2_ has not been previously described. To investigate this, we activated sorted T cells with an anti-CD3 antibody followed by TL1A, or vehicle, for 48 hours. Given the lack of commercial anti-LTβ antibodies, cell-surface detection of the functional heterotrimer (LTα_1_β_2_) was performed using a combination of an anti-LTα and a LTβR-Fc chimera (Supplementary Figure 3F). In agreement with our predictions, stimulation with TL1A induced a significant increase in LTα_1_β_2_ surface expression (Figure 3G). These results demonstrate a previously unrecognized TL1A/LTα_1_β_2_ axis and suggest its potential activation in the rectal mucosa of patients with PFD in the absence of rectal involvement.

### TL1A-LTα_1_β_2_ signaling as an alternative pathway in the context of TNF inhibition

TNF inhibitors are currently the only recommended biologic to treat active PFD in CD patients(16), suggesting the involvement of TNF in the development or maintenance of fistula tracts(17). Hence, we investigated whether TNF could also induce the PFD-associated signature. We first tested the effect of TNF on CD3+ peripheral T cells and observed that TNF increases *LTB* transcription (Figure 4A), in agreement with previous observations(18). However, in contrast to TL1A, TNF had no effect on the protein expression of the functional heterotrimer LTα_1_β_2_ (Figure 4B). Conversely, TNF, as observed with LTα_1_β_2_, significantly modulated the expression of the S1 CD+PFDun deregulated genes *TFPI2* and *CTGF*, as well as of the PFD-associated genes *MMP1* and *MMP3*, but not *CHI3L1*, in primary cultured fibroblasts (Figure 4C).

**Figure 4:**
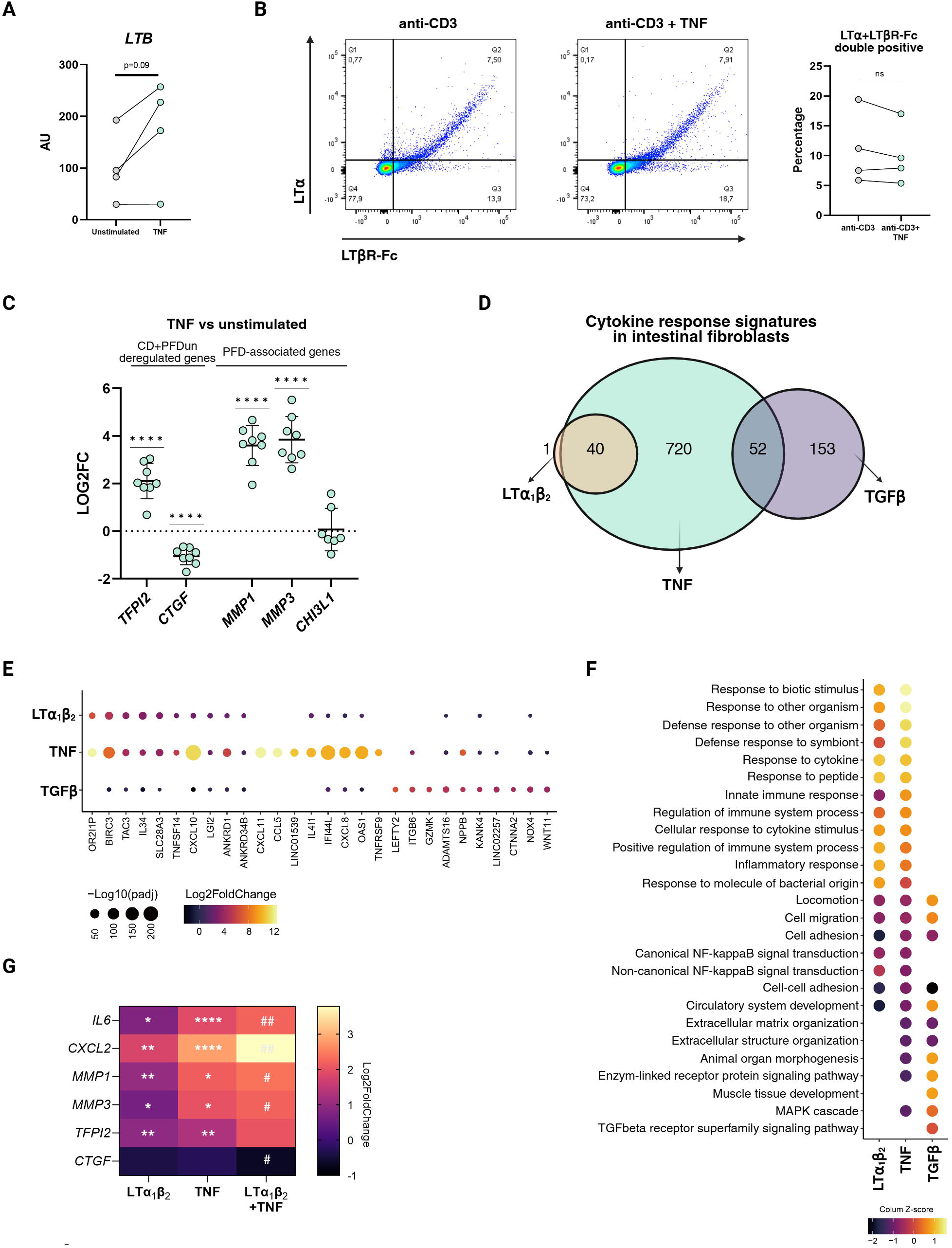
TNF induces the PFD-associated signature in primary fibroblasts. **A:** qRT-PCR for *LTB* in peripheral T cells unstimulated or stimulated with TNF for 48 hours. Paired t-Test, parametric, ** = p ≤ 0.01, n = 4. **B:** Flow cytometry plots of peripheral T cells stimulated with anti-CD3 or anti-CD3+TNF for 48 hours and stained with a-LTα and LTβR-FC. Double-positive cells correspond to LTα_1_β_2_+ T cells. Plots are from a representative replicate, and percentages of LTα_1_β_2_+ T cells comparing the two conditions are represented on the right. Paired t-Test, parametric, ns = p ≥ 0.05, n = 4. **C:** qRT-PCR of primary intestinal fibroblasts stimulated with TNF (20ng/ml) for 24 hours. Data is shown as a logarithm of the fold change in base 2 (Log2FC) comparing TNF-stimulated *vs* unstimulated fibroblasts. One sample t-Test, parametric, **** = p < 0.0001, n = 8. **D:** Venn diagram of the differentially expressed genes (DEGs) in the primary fibroblasts stimulated with LTα_1_β_2_ (orange), TNF (green) or TGFβ (purple). **E:** Dot plot with the top 10 genes induced in primary fibroblasts stimulated with LTα_1_β_2_, TNF and TGFβ. Dot size indicates adjusted p-value in the logarithm base 10, while color scale indicates the magnitude of Log2FC. **F:** Pathway analysis from the RNA sequencing data of the primary fibroblasts stimulated with LTα_1_β_2_, TNF and TGFβ. Color scale indicates adjusted p-value obtained from the enrichment test for biological pathway analysis by “ToppFun” in a logarithm base 10, scaled for each treatment. Data is represented as a Z-score. **G:** Heatmap representing the qRT-PCR data of primary intestinal fibroblasts stimulated with LTα_1_β_2_ (100ng/ml) and/or TNF (0.5ng/ml) for 24 hours. Data is shown as a logarithm of the fold change in base 2 (Log2FC) comparing stimulated *vs* unstimulated fibroblasts. LTα_1_β_2_ and TNF conditions: One sample t-Test, parametric, * = p < 0.05, ** = p < 0.01, *** = p < 0.001. TNF *vs* LTα1β_2_+ TNF condition: Paired t-Test, # = p < 0.05, ## = p < 0.01, n = 5.

To better characterize the response of intestinal fibroblasts to TNF and LTα_1_β_2_, we sequenced RNA from fibroblasts stimulated with each cytokine. Likewise, TGFβ-stimulated fibroblasts were also sequenced for signature comparison. A total of 1,276 genes were significantly regulated in response to TNF (822 up and 454 down), while 316 genes responded to TGFβ (205 up and 111 down) and 41 genes were significantly upregulated by LTα_1_β_2_ (Supplementary Table 4). Among the LTα_1_β_2_-induced genes, 40 genes (>97%) overlapped with the TNF-induced signature on fibroblasts (p=6.14e-55, Fisher’s test), while no overlap was found between LTα_1_β_2_ and TGFβ-induced signatures (Figure 4D). In contrast, a partial but significant overlap between TNF- and TGFβ-induced genes was observed (p=8.57e-27, Fisher’s test). Genes regulated by both TNF and LTα_1_β_2_ included NFκB subunits (*NFKB2*, *RELB*), genes related to interferon signaling (*OASL*) and immune responses (*IL34*, *CXCL10, VCAM1*) (Figure 4E and Supplementary Table 4). Conversely, sharing signatures between TNF and TGFβ were involved in ECM organization and MAPK cascade pathways (Figure 4F). Hence, while all three cytokines are fibroblast modulators, they each imprint a unique response, with TNF showing the most overlap with both LTα_1_β_2_, and TGFβ.

While our *in vitro* data shows that LTα_1_β_2_ and TNF could each independently induce the fibroblast signature associated with PFD, most patients in the CD+PFDun group (5/7) were under anti-TNF treatment at the time of inclusion (Supplementary Table 1), and none had rectal involvement (previous or current luminal disease). This suggests that the PFD signature in this group may be driven predominantly by TL1A/LTα_1_β_2_. Nonetheless, a role for TNF in promoting fistula formation cannot be ruled out in the context of intestinal inflammation. In fact, TNF, even at low concentrations (0.5ng/ml), synergizes with LTα_1_β_2_ to induce common target genes (i.e., *IL6, CXCL2*), including genes from the PFD signature (i.e., *MMP1* and *MMP3*) (Figure 4G). Altogether, our results suggest that the TNF and the TL1A/LTα_1_β_2_ axis exert independent and synergistic effects in promoting fibroblast activation in PFD.

### PFD is associated with an inflammatory-like signature in the rectal epithelium

The current hypothesis postulates that transitional cells lead to the formation of fistula tracks, as a result of epithelial cells undergoing EMT(5). Hence, we analyzed the activity of the EMT pathway in our scRNA-seq data, but found no evidence of activated EMT signature when comparing CDun and CD+PFDun patients (Supplementary Figure 4A). However, our analysis revealed a large number of DEGs between these patients, particularly within the undifferentiated transit amplifying (TA) cells and the top-most differentiated AQP8+ enterocytes (Supplementary Figure 1D, Supplementary Table 3). Pathway analysis showed the upregulation of inflammation-related processes, such as the defense response to bacterium and virus, and the response to cytokines such as type I interferon or IL-13 in AQP8+ enterocytes from CD+PFDun samples (Figure 5A). Moreover, we also observed an upregulation of processes related to oxidative phosphorylation, involving genes such as *COX5B*, *CYBA* and *DUOXA2*. Indeed, *DUOXA2,* the top fifth gene overexpressed in AQP8+ enterocytes from CD+PFDun patients, was also significantly upregulated in whole tissue rectal biopsies from those patients compared to the CDun group (Supplementary Figure 4B). Regarding TA cells, processes related to superoxide anion generation (*CYBA*, *COX5A*) and immune response (including the TL1A receptor *TNFRSF25*, *TNFRSF1A*, *CXCL2*) were also upregulated, along with a strong enrichment in processes involving the endoplasmic reticulum and the unfolded protein response (UPR) (*XBP1*), suggesting that these cells are undergoing high translational activity (Figure 5B). Interestingly, we also observed that the most upregulated (highest FC) gene in CD+PFDun patients was *OLFM4* (Figure 5B, C). Olfactomedin-4 is a glycoprotein involved in mucosal defense responses specifically produced by undifferentiated epithelial cells in the lower crypt, whose expression dramatically increases in the inflamed epithelium(19). We confirmed the upregulation of Olfactomedin-4 by protein staining (Figure 5D) and bulk RNA analysis of rectal biopsies (Supplementary Figure 4B) in the absence of previous or ongoing disease activity, suggesting inflammation-independent mechanisms are modulating the epithelial response in CD+PFDun patients.

**Figure 5:**
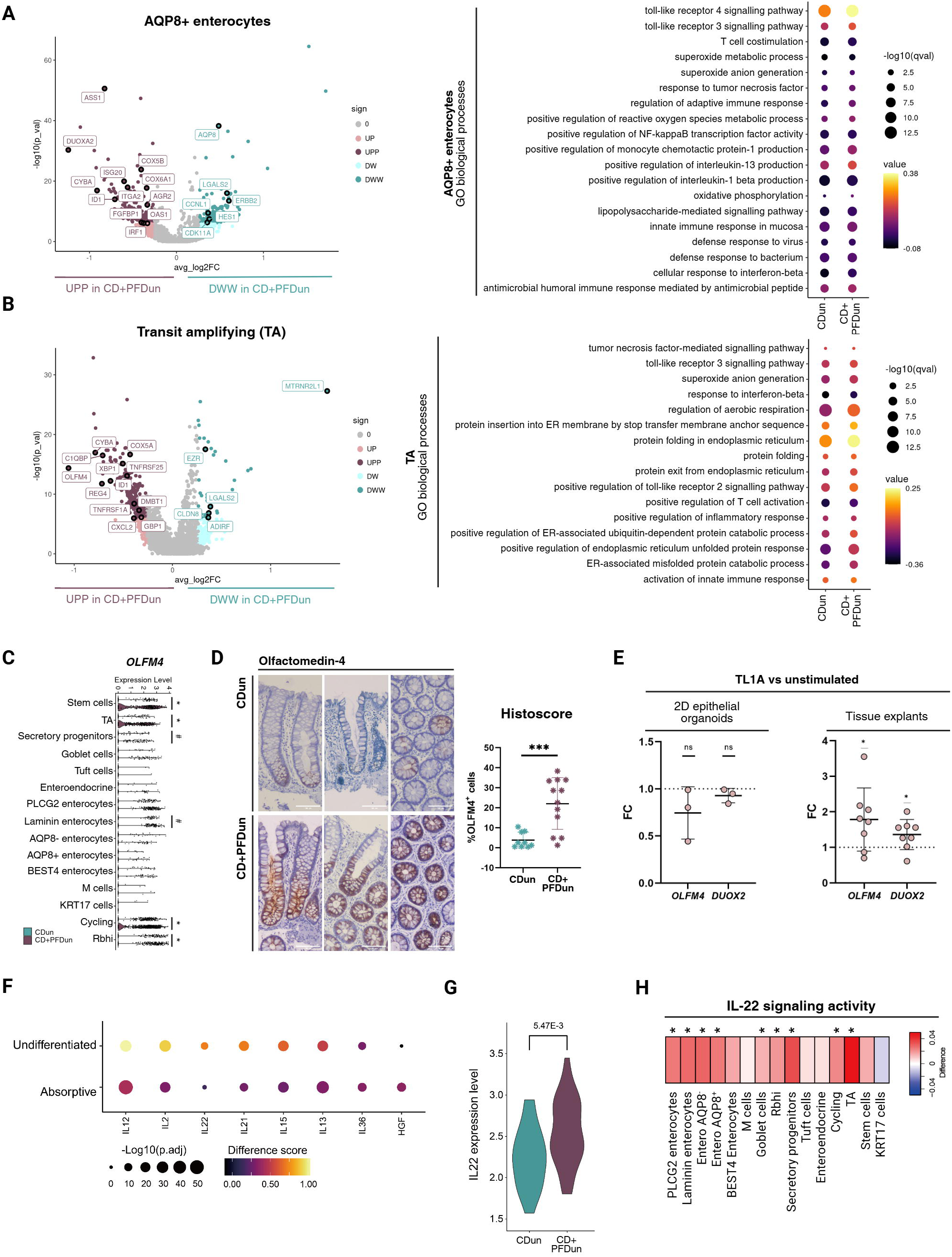
The epithelium of CD+PFDun patients presents activation downstream of IL-22 signaling. **A and B:** On the left, volcano plot representing the differentially expressed genes (DEGs) in the AQP8+ enterocytes (**A**) or the transit amplifying cells (TA) (**B**) between CDun and CD+PFDun patients. A two-sided Wilcoxon rank sum test was applied. Genes with an adjusted p-value < 0.05, and an avgLog2FC ˃ log2 (1.2) or avgLog2FC ˂ -log2 (1.2) were considered up-(UPP) or down-regulated (DWW), respectively. Genes with a nominal p-value ˂ 0.05 are also indicated in light colors if avgLog2FC ˃ log2 (1.2) (UP) or avgLog2FC ˂ -log2 (1.2) (DW). On the right, dot plot indicating pathways upregulated in the AQP8+ enterocytes (**A**) or TA (**B**) of CD+PFDun patients. Dot size indicates adjusted p-value in logarithm base 10, while color scale indicates the arbitrary value of each biological process generated by the AddModuleScore function in the Seurat package. The Wilcoxon-rank sum test (two-sided) with Holm correction was applied to compare CDun and CD+PFDun patients. **C:** Violin plot of *OLFM4* expression in epithelial cells. P-value was obtained by DEGs analysis. * = adjusted p-value < 0.05 and avgLog2FC ˃ log2 (1.2), # = nominal p-value ˂ 0.05 and avgLog2FC ˃ log2 (1.2). **D:** Olfactomedin 4 (OLFM4) immunohistochemistry staining of CDun and CD+PFDun formalin-fixed rectum biopsies. Representative images of tissue sections per each group are shown. Percentage of OLFM4+ cells were scored with the software QuPath. Unpaired t-Test, *** = p < 0.001. Scale bar: 100µm. **E:** qRT-PCR of organoid-derived monolayer (left) and cultured tissue explants (right) after 24 hours cultured with or without TL1A (300ng/ml). Data are represented as the fold change (FC) relative to the unstimulated condition. One sample t-Test, * = p < 0.05, ns = p > 0.05. **F:** Dot plot indicating top 5 cytokines explained by CytoSig for undifferentiated and absorptive cells. Dot size indicates adjusted p-value in logarithm base 10, while color scale indicates the difference in Z-score between CD+PFDun and CDun. **G:** Violin plot of *IL22* expression in the *IL22*+ expressing cell types (ILC3, Th17, CD4 effector memory and CD4 effector TNF). P-value was obtained by DEG analysis considering only *IL22* expressing cells. **H:** Heatmap indicating the difference score of IL-22 pathway activity by AddModuleScore of CD+PFDun relative to CDun patients. The Wilcoxon-rank sum test (two-sided) with Holm correction was applied to compare CDusn and CD+PFDun patients. Asterisks indicate significant differences (adjusted p-value < 0.05).

### Changes in PFD-associated epithelium are driven by TL1A-dependent IL-22

Increased expression of the receptor for TL1A (*TNFRSF25)* in TA epithelial cells (Figure 5B) suggested a potential effect of TL1A on the epithelium in CD+PFDun patients. To test this, we stimulated with TL1A intestinal 2D epithelial monolayers or tissue explants from non-inflamed rectal biopsies. While TL1A did not regulate *OLFM4* or *DUOXA2* expression on epithelial cultures, it significantly induced the expression of both epithelial genes in tissue explants (Figure 5E), suggesting that TL1A might drive its effect in the epithelium by acting on non-epithelial cells.

To investigate potential upstream regulators of the PFD-associated epithelial signature, we applied Cytosig to the epithelial compartment(20). Several cytokines were significantly predicted to drive the PFD epithelial signature, including IL-12, IL-2, and IL-13 on absorptive cells, and IL-12, IL-2, and IL-22 on undifferentiated cells (Figure 5F). Among them, IL-13, IL-2, and IL-22 have previously been described to be induced by TL1A(21–24), but only the receptors for IL-13 and IL-22 were expressed on epithelial cells (Supplementary Figure 4C). Hence, we looked for differences between groups in the expression of these cytokines in our dataset. While no differences were detected for *IL13* (Supplementary Figure 4D), we observed that CD+PFDun patients expressed higher levels of *IL22* in IL-22-producing cell types (ILC3, Th17, CD4 effector memory and CD4 effector TNF, Supplementary Figure 4E) compared to CDun individuals (Figure 5G). Mapping the epithelial-specific IL-22 signature from an in-house microarray dataset to our scRNA-seq confirmed that this pathway was more active in most epithelial cell types from CD+PFDun patients, and it was significantly enriched in 10 out of 15 cell types (Figure 5H). This was further confirmed in a 2D-monolayer epithelial culture, where IL-22 induced the significant overexpression of *DUOX2* and *OLFM4*, along with other genes upregulated in CD+PFDun patients (Supplementary Figure 4F). Moreover, when we measured IL-22 protein production in the supernatants of the TL1A-stimulated tissue explants, we observed a significant correlation between *DUOX2* and IL-22 levels, although this did not reach significance for *OLFM4* (Supplementary Figure 4G). Finally, we confirmed that TL1A, but not TNF, could induce the secretion of IL-22 by anti-CD3-stimulated peripheral T cells (Supplementary Figure 4H). Overall, these results suggest that increased TL1A signaling in patients with PFD drives increased levels of IL-22, which in turn can induced an anti-bacterial response on the epithelial compartment of these patients.

## DISCUSSION

PFD in CD remains a significant clinical challenge with poorly understood pathophysiology. Most previous studies have focused on fistula or peri-fistula tissues to identify potential therapeutic interventions aimed at treating already established fistulizing disease. In contrast, few have investigated the predisposing molecular mechanisms that drive fistula development. In that sense, several genetic variants have been found that increase the risk of perianal disease in CD patients(25), supporting the existence of patient-specific predisposing factors. Beyond genetic predisposition, to identify novel mechanisms of fistula formation we examined tissues distant from the fistula tract but within a relevant anatomical context (i.e., rectal mucosa) of CD patients with and without PFD, and used an unbiased approach that relied on whole-transcriptome sequencing of all rectal mucosa cells.

Using this strategy, we identified changes in cell abundances and in transcriptional signatures across cell types, with the most pronounced alterations observed in lamina propria fibroblasts, T cells and epithelial cells. The fibroblast signature associated with the perianal phenotype was at odds with the classical view that implicated TGFβ in the pathophysiology of PFD(4,5,26), but was in line with recent studies using high throughput analyses of fistula tracts, which did not report any higher activity in TGFβ signaling either(27,28). In contrast, our data supports the role of TNF family members (LTα_1_β_2_ and TNF) in modulating the transcriptional states of fibroblasts in the lamina propria of patients with PFD. While the role TNF plays in driving fistula development(17) and treating established tracts(16) has been well described, no previous data had addressed LTα_1_β_2_ in the pathophysiology of PFD.

Lymphotoxin β (*LTB*), which we found to be significantly overexpressed in mucosal lymphocytes from patients with PFD, forms a surface-bound heterotrimer with lymphotoxin α (LTα_1_β_2_), which exclusively binds to the lymphotoxin β receptor (*LTBR*) expressed by stromal, epithelial and some myeloid cells(29). Largely known for its role in organizing and maintaining the structures of lymphoid tissues, LTα_1_β_2_-LTβR signaling is also involved in initiating immune responses, such as those protecting against microbial pathogens in mucosal barriers(30,31). Based on our scRNA-seq data, all mesenchymal cell types express *LTBR*, being the interaction between *LTB-LTBR* predicted to be more predominant in CD+PFDun patients. In line with its known function, LTα_1_β_2_ induced genes associated with the organization of the lymphoid follicle and with immune responses (*CXCL2*, *IL6* or *VCAM1*), significantly overlapping with the TNF-induced gene signature. Notably, LTα_1_β_2_ also induced some of the genes overexpressed in the lamina propria S1 fibroblasts of CD+PFDun patients, as well as *MMP1*, *MMP3* and *CHI3L1*, supporting a role for LTα_1_β_2_-LTβR signaling in PFD. Interestingly, a preprint from Constable *et al.*(28) reports that while fistula tissue from CD patients and patients with cryptoglandular disease were transcriptionally similar, *LTB* and *LTA* were two of the few genes overexpressed in CD, strengthening our hypothesis. While our data also supports a role for TNF in imprinting the stroma of PFD patients, TL1A, but not TNF, was identified as the upstream inducer of LTα_1_β_2_ expression on activated T cells, thus revealing a novel therapeutic target independent of TNF(33). Despite TL1A (*TNFSF15*) not being increased in patients with PFD, the immune response enrichment analysis of CD4+ T cells, as well as the overexpression of both *LTB* and *IL22*, strongly reflects higher downstream TL1A activity in CD+PFDun patients.

TL1A is currently under investigation for its potential therapeutic value in IBD(32). A role for TL1A has been proposed as both an inflammation promoting agent and as a mediator of fibrostenosis based on preclinical studies(33). The potential involvement of TL1A in complicated phenotypes is supported by genome-wide association studies describing susceptible loci for *TNFSF15* in European and Japanese cohorts(34,35). In a Korean cohort, another non-risk single nucleotide polymorphism (SNP) allele of *TNFSF15 (*rs4574921) was also associated with a higher cumulative risk for developing PFD(36). However, to our knowledge, no previous study had functionally involved TL1A in the pathophysiology of PFD.

Beyond its novel effects on LTα_1_β_2_ production by CD4+ T cells, we also confirmed the ability of TL1A to induce IL-22 expression by lymphoid cells(21,22). A role for IL-22 in established PFD has recently been suggested, including a role in tissue remodeling(6,28) and EMT(6). While we found no evidence of EMT activation in the rectal epithelium of CD+PFDun patients, immune and anti-bacterial responses, including IL-22 targets, were upregulated in these patients, despite the non-involved and non-inflamed nature of the tissue. The overall epithelial response, imprinted by IL-22, suggests an increase in bacteria recognition, which would be supported by recent data showing an increased detection of serum antibodies against bacterial antigens such as anti-ASCA, anti-OmpC or anti-flagellins in CD patients with PFD(25,37).

While our study represents a step towards identifying the molecular mechanisms driving the development of perianal fistulae in CD, it has some limitations. First, the number of subjects per study group is limited and a validation cohort will be required. However, the TL1A-LTα_1_β_2_/IL-22 axis was validated in primary intestinal cultures. Most importantly, regardless of the upstream drivers (TL1A/LTα_1_β_2_ or TNF), the functional implications of the PFD-associated signature remain unknown. Based on our analysis, we propose a potential metabolic reprogramming of CD+PFDun lamina propria fibroblasts, with an enrichment in oxidative phosphorylation and lipid metabolism processes. Data from fibrotic skin fibroblasts shows that a metabolic switch from fatty acid oxidation to glycolysis can promote ECM accumulation(38). Likewise, it has been observed that fibroblasts can also sense mechano-physical alterations in the ECM, as a softer matrix promotes cholesterol and neutral lipid synthesis, whereas stiff ECM sustains glycolysis(39). This suggests that the rectum of CD+PFDun patients might present a less stiff ECM, which could be sensed and sustained by lamina propria S1 fibroblasts. This hypothesis is consistent with the upregulation of *MMP1* and *MMP3* in CD+PFDun patients, involved in matrix degradation. Nonetheless, further studies are required to demonstrate a direct link between fibroblast metabolic changes and fistula development.

To conclude, our data provides novel insights into the molecular mechanisms underlying the development of PFD and points to TL1A as a key upstream regulator in this complication. While our results also support a role for TNF in this phenotype, we identified a new signaling axis independent of, and potentially complementary to, TNF. This opens new questions, including how the changes we uncovered lead to the formation of fistulous tracts and, more importantly, whether interfering with TL1A signaling could prevent this process.

## ACKNOWLEDGMENTS

We thank the Single Cell Genomics Group at CNAG (Barcelona) and the Cytometry Core Facility of the IDIBAPS for their technical help.

## ABBREVIATIONS

DEGs: Differential expression genes
ECM: Extracellular matrix
ELISA: Enzyme-linked immunosorbent assay
EMT: Epithelial-to-mesenchymal transition
FFPE: Formalin and paraffin-embedded
IER: Immediate-early response
LTα_1_β_2_: Lymphotoxin alpha 1 beta 2
MMPs: Matrix metalloproteinases
PBMCs: Peripheral blood mononuclear cells
PFD: Perianal fistulizing disease
qPCR: Real-time PCR
scRNA-seq: Single-cell RNA sequencing
SNPs: Single nucleotide polymorphism
TGFβ: Transforming growth factor beta
TL1A: Tumor necrosis factor-like ligand 1A
TNF: Tumor necrosis factor
UPR: Unfolded protein response

## AUTHOR’S CONTRIBUTION

VG: Data acquisition and curation, methodology, investigation, formal analysis, visualization, writing – original draft; JWC: Data curation, formal analysis, software, visualization, writing – review and editing; BC: Investigation, resources, visualization. AMC: Data curation, software; MV: Methodology, investigation, formal analysis, writing – review and editing; ID: Methodology, investigation, formal analysis, writing – review and editing; LMG: Investigation, resources; AS-M: Data curation, software; EM-A: Methodology, investigation, writing – review and editing; MCM: Investigation, resources; ME: Investigation; IT: Investigation; LS: Investigation; AG: Investigation, resources; CP: Investigation, resources; EF: Investigation; RF-L: Investigation, resources, writing – review and editing; AM-C: Investigation, resources; CL: Investigation, resources; ME: Investigation, resources, writing – review and editing; JR: Investigation, resources; AF-C: Investigation, resources; IO: Investigation, resources; ER: Investigation, resources, writing – review and editing; JP: Conceptualization, investigation, funding acquisition, writing – review and editing; MH: Formal analysis, supervision, writing – review and editing; AS: Conceptualization, data curation, formal analysis, supervision, project administration, funding acquisition, writing – original draft.

## DATA TRANSPARENCY STATEMENT

The raw data generated in this work will be publicly available upon acceptance of the manuscript in the Gene Expression Omnibus (GEO).

## TABLE LEGENDS

Supplementary Table 1: Patients clinical data

Supplementary Table 2: Cell annotation

Supplementary Table 3: DEGs scRNA-seq data of CDun vs CD+PFDun patients

Supplementary Table 4: DEGs of TNF-, LTα_1_β_2_- and TGFβ-stimulated fibroblasts Supplementary

Supplementary Table 5: TaqMan primer probe references

## SUPPLEMENTARY FIGURE LEGENDS

**Supplementary Figure 1:**
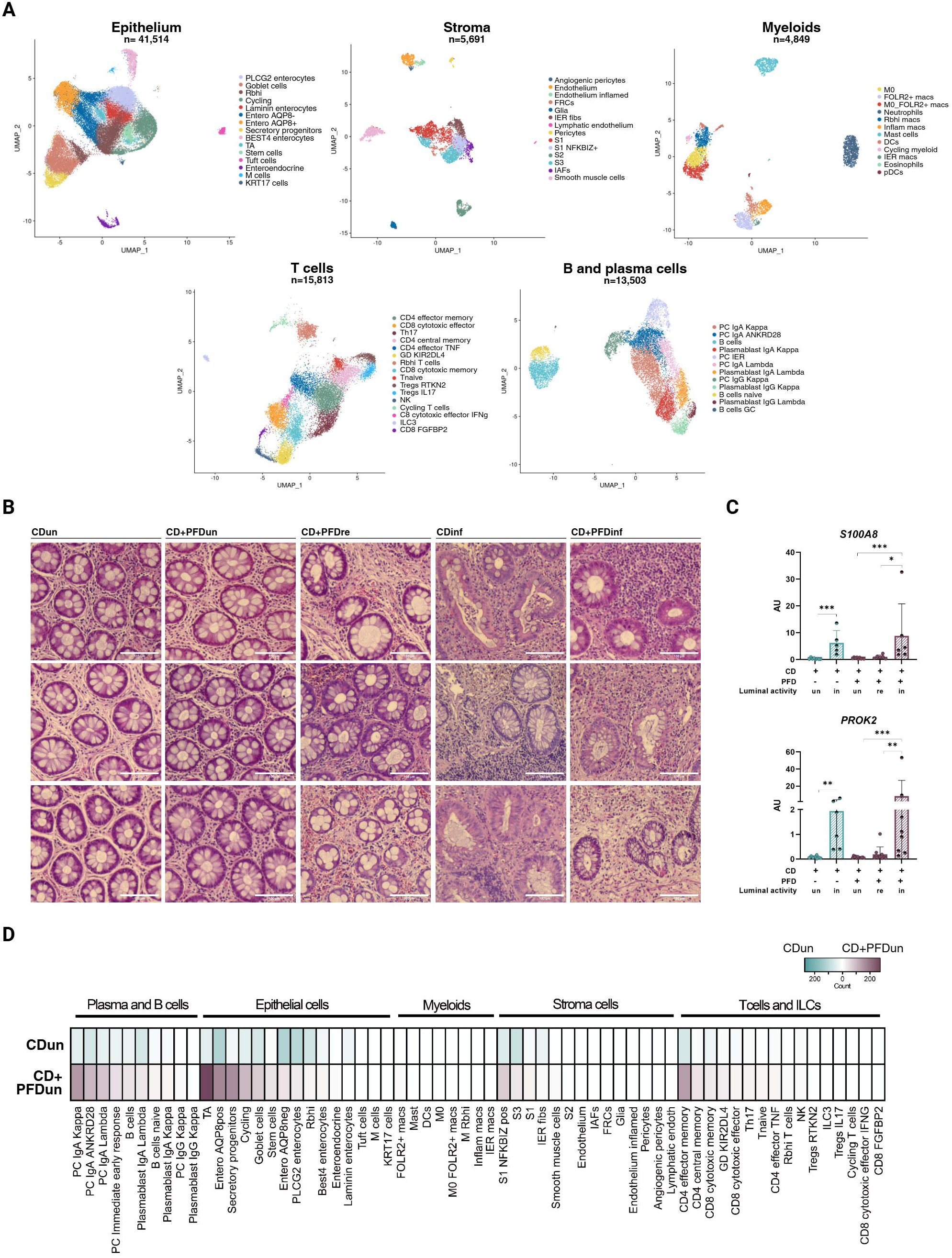
Transcriptomic differences between CDun and CD+PFDun patients are not driven by luminal inflammation. **A:** Uniform Manifold Approximation and Projection (UMAP) representation of the scRNA-seq dataset divided into the five main cell types. The number of cells analyzed in each subset is indicated. **B:** Hematoxylin and eosin staining of the rectal mucosa of patients from the cohort. Three representative images from each group are shown. Scale bar: 100µm. **C:** Whole-tissue qRT-PCR analysis of *S100A8* and *PROK2* expression in rectal biopsies from CDun (n=8), CDinf (n=6), CD+PFDun (n=12), CD+PFDre (n=10) and CD+PFDinf (n=8). AU: arbitrary units. Kruskal-Wallis test, * = q **<** 0.05; ** = q < 0.01, *** = q < 0.001. **D:** Heatmap representing the number of differentially expressed genes (DEGs) when comparing the cell types indicated between CDun and CD+PFDun patients. Only cell types with cell counts above 10 are shown.

**Supplementary Figure 2:**
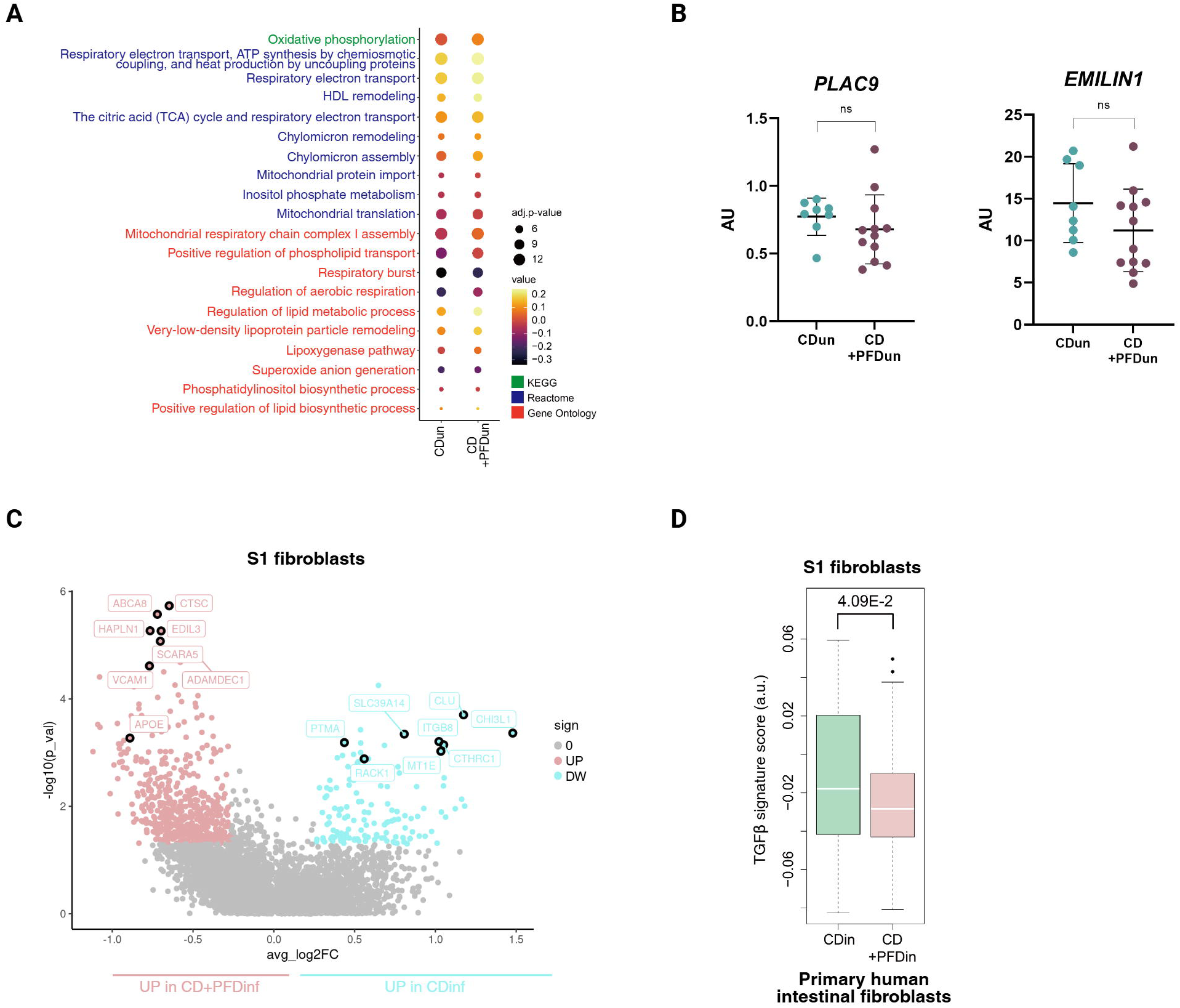
Alterations in the S1 fibroblasts of CD+PFDun patients occur independently of transforming growth factor beta (TGFβ). **A:** Dot plot indicating pathways upregulated in the S1 fibroblasts of CD+PFDun compared to CDun patients related to calcium and lipid metabolism. Dot size indicates adjusted p-value in logarithm base 10, while color scale indicates the arbitrary value of each biological process generated by the AddModuleScore function in the Seurat package. The color of the processes indicates the database they are derived from (KEGG, Reactome or Gene ontology). The Wilcoxon-rank sum test (two-sided) with Holm correction was applied to compare CDun and CD+PFDun patients. **B:** Whole-tissue qRT-PCR analysis of *PLAC9* and *EMILIN1* expression in rectal biopsies from CDun (n=8) and CD+PFDun (n=12), Mann-Whitney test, ns = p ≥ 0.05. **C:** Volcano plot representing the differentially expressed genes (DEGs) in the lamina propria S1 fibroblasts between CDinf and CD+PFDinf patients. Genes with a nominal p-value ˂ 0.05 are indicated, if avgLog2FC ˃ log2(1.2) (UP) or avgLog2FC ˂ -log2(1.2) (DW). **D:** Barplot indicating the TGFβ signature scores of S1 fibroblasts from CDinf and CD+PFDinf patients generated from bulk RNA sequencing data of TGFβ-stimulated primary fibroblasts mapped to the single-cell dataset. The Wilcoxon-rank sum test was applied. AU = arbitrary units.

**Supplementary Figure 3:**
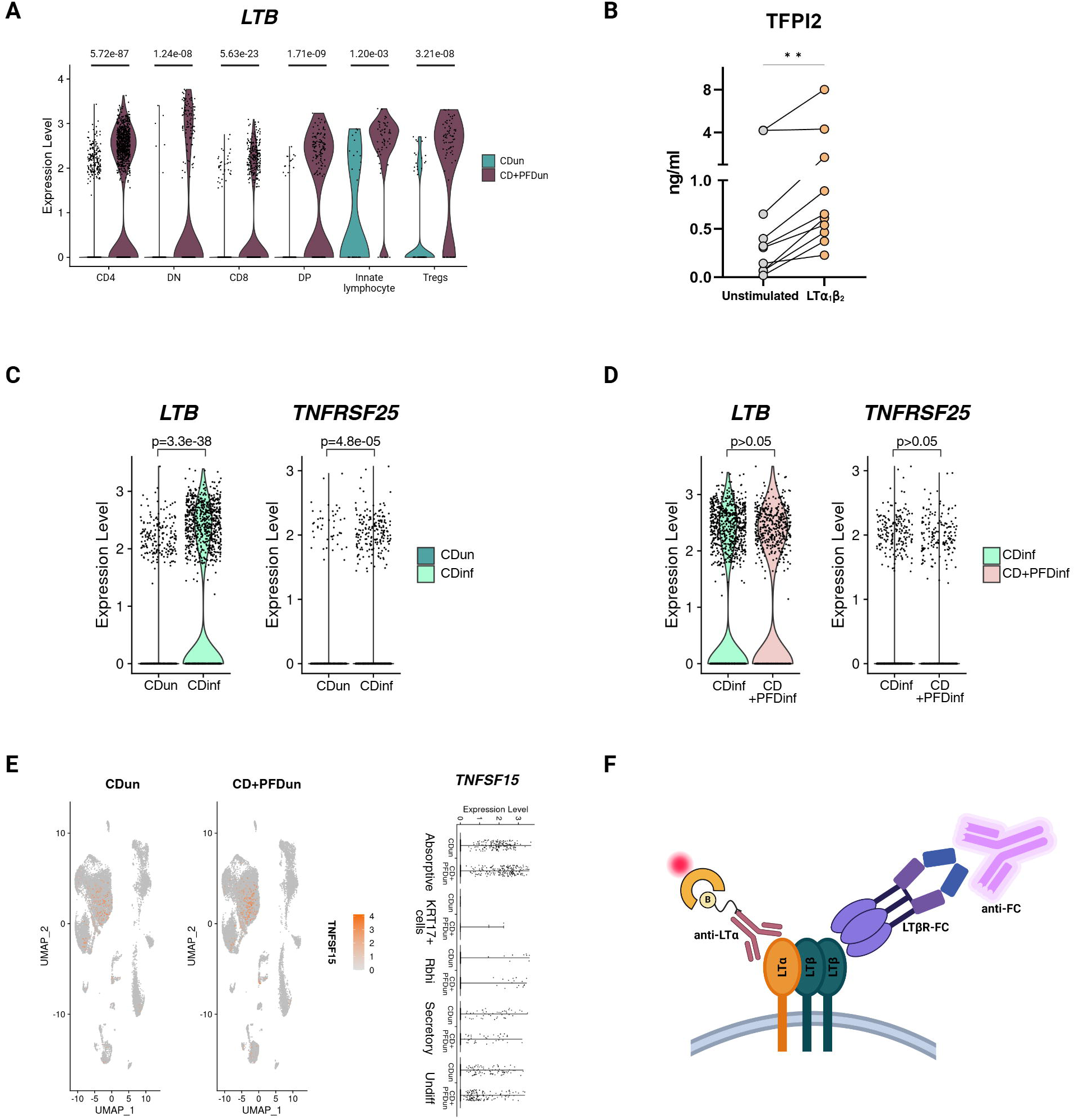
TL1A induces higher expression of lymphotoxin β in the T cells. **A:** Violin plot for *LTB* expression in the T-cell compartment. The adjusted p-values obtained by DEG analysis are indicated for each comparison. **B:** ELISA for TFPI2 in the supernatants of primary intestinal fibroblasts stimulated with LTα_1_β_2_ (100ng/ml) for 24 hours compared to the unstimulated condition. Paired T test, nonparametric, n=10, ** = p < 0.01. **C:** Violin plot for *LTB* and *TNFRSF25* expression in the CD4+ T cell compartment, comparing CDun and CDinf patients. The adjusted p-values generated by DEG analysis are indicated. **D:** Violin plot for *LTB* and *TNFRSF25* expression in the CD4+ T cell compartment, comparing CDinf and CD+PFDinf patients. The adjusted p-values generated by DEG analysis are indicated. **E:** On the left, Uniform Manifold Approximation and Projection (UMAP) plots of the whole object showing the expression of *TNFSF15*. On the right, violin plots indicating the expression of *TNFSF15*. **F:** Graphical representation of the flow cytometry approach to detect LTα_1_β_2_+ T cells.

**Supplementary Figure 4:**
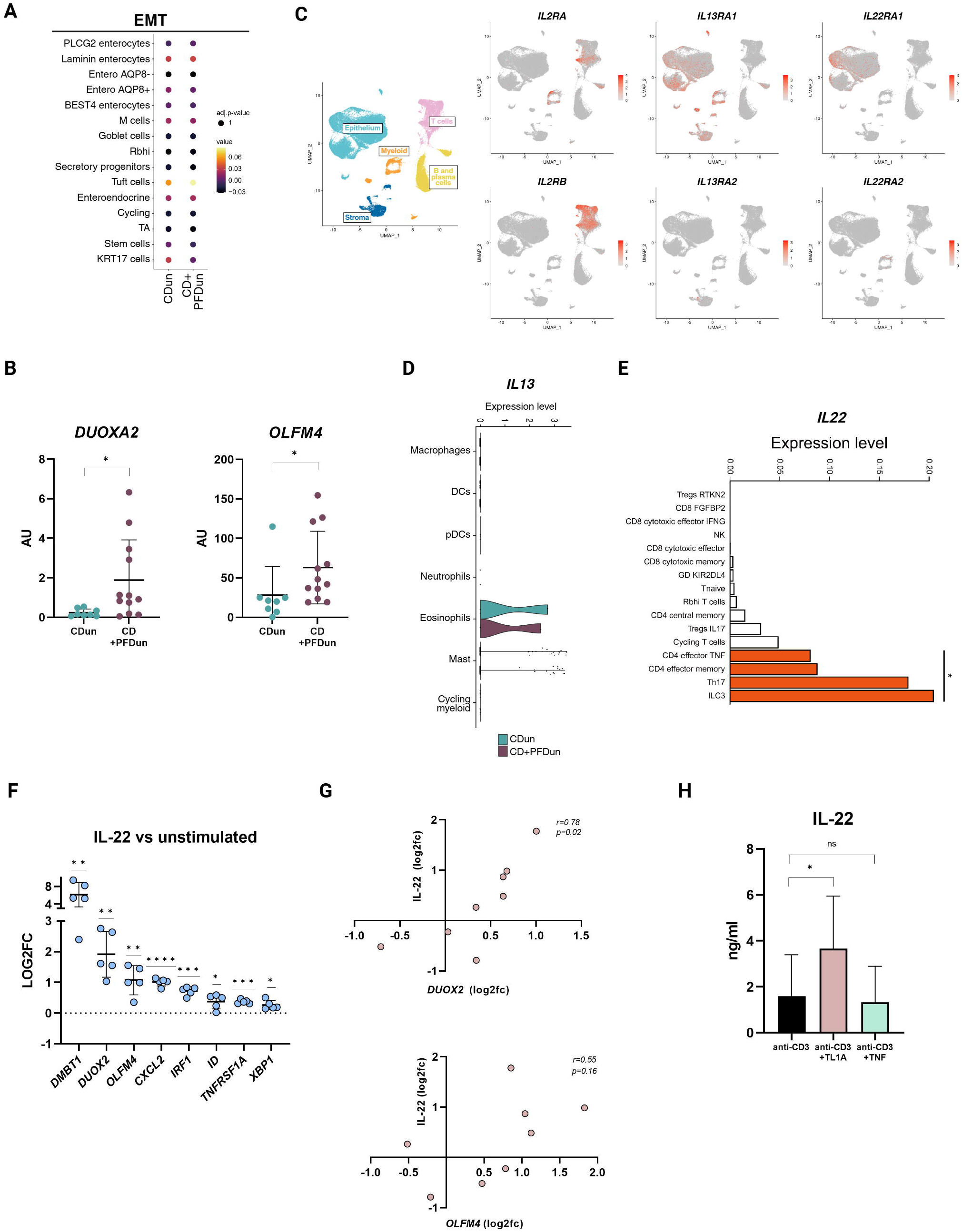
The epithelium of CD+PFDun patients present activation downstream of IL-22 signaling. **A:** Dot plot representing the activity of the “epithelial-to-mesenchymal transition” (EMT) process (Gene Ontology) in each epithelial cell type comparing CDun and CD+PFDun patients. Dot size indicates adjusted p-value, while color scale indicates the arbitrary value of the pathway generated by the AddModuleScore function in the Seurat package. The Wilcoxon-rank sum test (two-sided) with Holm correction was applied to compare CDun and CD+PFDun patients. **B:** Whole-tissue qRT-PCR analysis of *DUOXA2* and *OLFM4* expression in rectal biopsies from CDun (n=8) and CD+PFDun (n=12) patients; Mann-Whitney test, * = q < 0.05, AU: arbitrary units. **C:** Uniform Manifold Approximation and Projection (UMAP) plots of the whole object, showing the expression of *IL2RA*, *IL2RB*, *IL13RA1*, *IL13RA2*, *IL22RA1*, and *IL22RA2*. **D:** Violin plot of *IL13* expression in myeloid cells. None of the comparisons were significant by DEG analysis. **E:** Bar plot indicating the expression of *IL22* in each of the T-cell subtypes. Bars in red indicate those cell types whose *IL22* expression was significantly higher than the rest. Adjusted p-value was obtained by FindAllMarkers function. * = adjusted p-value < 0.05. **F:** qRT-PCR of epithelial organoids seeded as a 2D monolayer upon stimulation with IL-22 (5ng/ml) for 24 hours. Data is represented as the fold change (FC) of stimulated organoids relative to the unstimulated condition. One sample T test, parametric, * = p < 0.05, ** = p < 0.01, *** = p < 0.001, **** = p < 0.0001. n= 5. **G:** Correlation analysis of *DUOX2* and *OLFM4* expression to the levels of IL-22 protein production in the tissue explants stimulated with TL1A for 24 hours. Data is represented as the logarithm in base 2 of the FC (log2fc) relative to the unstimulated condition. *r* and p-values are indicated. **H:** Protein production of IL-22 in the supernatants of peripheric CD3+ T cells cultured for 48h and stimulated with anti-CD3, anti-CD3+TL1A or anti-CD3+TNF. Paired T test analysis compared to the anti-CD3 condition, * = p < 0.05, ns = p > 0.05.

## SUPPLEMENTARY INFORMATION

### Patient recruitment and sample collection

Rectal biopsies (n=8 from each patient) from CD patients with or without perianal fistulizing disease (PFD) were collected to perform single-cell RNA sequencing (scRNA-seq), bulk RNA analysis and tissue staining analysis. For histology analysis, biopsies were fixed in formalin and paraffin-embedded (FFPE). Additionally, rectal biopsies were collected for tissue explant cultures. Biopsies were collected during routine endoscopies performed as standard care at the Hospital Clinic (Barcelona, Spain) or at the Hospital Universitari Mutua Terrassa (Terrassa, Spain) upon signature of an IRB-approved informed consent. Primary fibroblast and epithelial organoid cultures were generated from the non-tumoral areas of colorectal cancer patients undergoing surgery (herein referred to as non-IBD patients). Buffy coats from healthy control patients (Banc de Sang i Teixits, Barcelona, Spain) were used to isolate CD3+ peripheral T cells.

### Ethics statement

The study was approved by the Ethics Committee of Hospital Clinic Barcelona (HCB/2020/0035 and HCB/2016/0389) and the Hospital Universitari Mutua Terrassa (CI201901). Participants received no compensation for the study.

### Patient and Public Involvement

Patients or members of the public were not involved in the design, conduct, reporting, or dissemination plans of this research.

### Tissue digestion for single-cell RNA-seq analysis

Rectal biopsies (n=4 per patient) were processed within 1 hour of collection. Tissue was digested as previously described(1). Briefly, biopsies were mechanically and enzymatically digested using 0.5 Wünsch units/ml Liberase^TM^ (Roche) and 10µg/ml DNAse I (Sigma) for 1 hour at 37°C and 250rpm agitation. After digestion, the cell suspension was filtered to obtain clean single-cell suspensions. Cells were counted and transferred to the single-cell genomics platform.

### Single-cell RNA sequencing of intestinal cells

Cell suspensions were processed using the 10× Genomics 3′ mRNA single-cell protocol with the Chromium System (10x Genomics, CA, USA). Around 7,000 cells were loaded into the Chromium platform following the manufacturer’s guidelines. The creation of gel beads in emulsion (GEMs), along with barcoding and reverse transcription within the GEMs, was carried out using the Chromium Single Cell 3′ and V(D)J Reagent Kits (10× Genomics, CA, USA), following the user’s manual (no. CG000086). The resulting full-length, barcoded cDNA was amplified via PCR using Nextera® PCR primers (Illumina, CA, USA) to obtain sufficient material for library preparation. Library sequencing was conducted on the HiSeq2500 platform (Illumina, CA, USA).

### Single-cell data processing

Sequences obtained in fastq files were processed with the CellRanger’s count pipeline with default parameters for each sample (10X Genomics Cell Ranger v3.1.0). This pipeline performs an alignment based on the reference genome (Gencode release 27, assembly GRCh38 p10), demultiplexing, and UMI counting. The resulting matrix was analyzed using R (v.4.1.2). First, we removed potential doublets using the scDblFinder (v1.2.0) function of the Celda R package(2). Next, we followed the default pipeline from the Seurat package (v4.0.3) for preprocessing, including bad cell removal, normalization, scaling, PCA, UMAP, and Louvain clustering(3). In detail, cells that expressed fewer than 200 unique genes or had a high-count ratio aligned to the mitochondrial transcriptome were removed. We also removed putative red blood cells that expressed *HBA1*, *HBA2*, and *HBB*. We then selected highly variable genes across the different samples. We selected the consensus top 1,500 genes across samples with random sampling for ties(4,5). Further removal of ribosomal proteins, mitochondrial genes, HLA genes, IG genes, and cell cycle-related genes was done to avoid batch effects(4–6). The “RunUMAP” function in Seurat v4 was utilized for dimensionality reduction. The neighbor graph for cells was obtained by “FindNeighbors,” which conducts shared-nearest-neighbor. Further Louvain clustering was conducted via the “FindClusters” function. As a result, we could make primary classifications (epithelial cells, stromal cells, T cells, plasma and B cells, and myeloid cells) before splitting cells for in-depth analysis. We excluded clusters showing doublet behavior. In addition, IG genes were removed from all the main cell types, except B cells, to reduce background noise.

### Batch correction

We ran Harmony (version 0.1.0) to correct the batch effect between every patient for each main cell type(7). After batch correction, we ran another round of preprocessing for each main cell type, except for giving the reduction parameter as “harmony.”

### Identification of cell types

We used marker gene expression to categorize cell clusters for the main cell types: *EPCAM*, *KRT8*, *KRT18*, *PYY*, and *SH2D6* for epithelial cells; *CD79A*, *DERL3*, and *MS4A1* for plasma and B cells; *CD3D*, *CD3E*, and *KLRB1* for T cells and ILCs; *PECAM1*, *KDR*, *CDH19*, *COL3A1*, *LUM*, and *MMP2* for stromal cells; *TPSAB1* and *CPA3* for mast cells; and *CD14* and *CD68* for other myeloid cells. The annotations for each subcluster from the main cell type were defined by the marker genes, obtained by the “FindAllMarkers” function with the default threshold, except for the min.pct parameter, which was set to 0.25. We further filtered out the marker genes by q values higher than 0.01. Detailed marker gene lists can be found in Supplementary Table 2.

### Cell abundance comparisons

The relative abundance of each cell type was measured by normalizing the total number of cell counts in each group. We performed Fisher’s exact test (one-sided; greater) between the two groups to obtain a p-value. Benjamini-Hochberg (BH) correction was conducted to obtain adjusted p values. We defined |FC| > 1.5 and adjusted p-value < 0.05 as a significant difference.

### Differential expression gene (DEG) analysis

The DEG analysis to compare the two groups was conducted using the “FindMarkers” function (logfc.threshold = 0.01 with default parameters) in the Seurat package(3). DEGs were defined by |avgLog2FC| > log2(1.2) and adjusted p-value < 0.05.

### Gene signature analysis

We measured the signature score by the “AddModuleScore” function in the Seurat package(3) using the Gene ontology(8,9) (2022-07-01, doi:10.5281/zenodo.6799722) database. We tested only with the overlapping genes between the database and those in our data with the “ctrl” parameter set to 5. We used the average score from the results of “AddModuleScore” for each cell. While comparing the two groups, we performed the Wilcoxon-rank sum test (two-sided) to obtain p values. Holm correction was conducted to obtain adjusted p values.

### Cell-cell interaction analysis

We used CellPhoneDB(10) (v2.1.7) to analyze the receptor–ligand-mediated cell-cell interactions. We used the raw count matrix with only the consensus coding gene sequence (CCDS)(11) overlapping genes based on the scRNA-seq data. We ran each group with the default pipeline and parameters. The strength of interaction was defined by “mean_exp” obtained from the CellphoneDB output.

### Identifying expressing cell types

We identified certain cell type(s) if they expressed a given factor to a significantly higher degree than the other cells by FindAllMarkers as described above with adjusted p-value < 0.01.

### Peripheral T-cell isolation and TL1A stimulation

Human peripheral blood mononuclear cells (PBMCs) were isolated from buffy coats of healthy controls with a Lymphoprep gradient (Nycomed Pharma) following the manufacturer’s instructions. After isolation, 40M of PBMCs were stained for CD3-APC (UCHT1 clone, 300412, BioLegend) and the Zombie Aqua Fixable Viability Kit (4231101, BioLegend) and sorted with FACS Aria II (BD Bioscience). One million sorted CD3+ T cells were seeded in 24-well plates pre-coated with 1µg/ml anti-CD3 (HIT3a clone, 300331 BioLegend) for 48 hours in triplicate, with or without 300ng/ml of TL1A (1319-TL, R&D). After incubation, cells were collected and centrifuged at 500*g* for 10min at 4°C and the pellet was either resuspended in 500µl of TRIzol (ThermoFisher) for RNA isolation or used for flow cytometry staining. Supernatant was collected for protein secretion analysis.

### Flow cytometry staining

CD3+ peripheric T cells were stained with biotinylated anti-LTα (BAF211, R&D) and LTβR-FC (7538-LR, R&D), and detected with streptavidin-PE (554061, BD Bioscience) and anti-human IgG-BV421 (109-675-170, Jackson ImmunoResearch), respectively. Viability was measured with the Zombie Aqua Fixable Viability Kit (423101, BioLegend). Cells were fixed using BD Stabilizing Fixative [BD], acquired using a BD FACSCanto II flow cytometer (BD), and analyzed with FlowJO software (version 10.6.1, BD).

### Isolation and culture of intestinal fibroblasts

Fibroblasts were isolated as previously described(12,13). Briefly, surgical resections from non-IBD patients were digested with agitation at 37°C with 1mM DTT and 5mM EDTA for 30min, followed by 30min-digestion with 5mM EDTA. The tissue was then mechanically and enzymatically digested with 5.4U/ml Collagenase D (Roche), 39.6 U/ML DispaseII (Gibco) and 100 U/ML DNAse I (Sigma) for 20min at 37°C and 250rpm agitation. Cell suspension was filtered through a 70µm cell strainer and centrifuged at 500*g* for 5min at RT. The pellet was seeded onto a T-25 flask and fibroblasts were selected by removing non-adherent cells. Cells were grown until they reached confluency and passaged using 0.25% Trypsin-EDTA (Gibco). For stimulation experiments, 40,000 fibroblasts per condition were seeded in a 24-well plate. After 3 days, fibroblasts were stimulated with 5ng/ml TGFβ (PHG9201, Invitrogen), 100ng/ml LTα_1_β_2_ (8884-LY, R&D) or 20ng/ml TNF (10291-TA, R&D) for 24 hours. After incubation, supernatant was collected, cells were washed with PBS (Gibco) and 500µl of TRIzol reagent (ThermoFisher) per condition was added to the well for RNA extraction.

### Epithelial isolation and 3D organoid cultures

Generation of epithelial organoids and establishment of 3D organoid cultures were conducted as previously described(14). In brief, epithelial crypts were isolated from non-IBD surgical samples after incubation with 8mM for 45min at 4°C. Isolated crypts were embedded in Matrigel (Corning) and cultured in organoid growth medium (Wnt3a-conditioned medium and Advanced DMEM/F12 50:50, 1x GlutaMAX 10 mM HEPES, N-2 (1×) (Gibco), B-27 without retinoic acid (1×) (Gibco), 10 mM nicotinamide (Sigma), 1 mM N-Acetyl-L cysteine (Sigma), 500 ng/mL R-spondin-1 (RSPO1, Sino Biologicals), 50 ng/mL human epidermal growth factor (EGF) (Invitrogen), 100 ng/mL human Noggin (Peprotech, Rocky Hill, NJ), 10nM Human [Leu15]-Gastrin I (Sigma), 500 nM LY2157299 (Axon MedChem, Groningen, The Netherlands), 10 µM SB202190 (Sigma), and 100nM prostaglandin E2 (Sigma). Organoids were passaged every 5-6 days using a dispase-based cell dissociation protocol(15). After a maximum of 3 passages, organoids were stimulated with 20ng/ml IL-22 (200-22, Peprotech) for 24 hours and harvested with TRIzol reagent (ThermoFisher) for microarray analysis.

### Organoid-derived 2D monolayer cultures

Organoids were dissociated to single cells to form 2D epithelial monolayers. Single cells were seeded onto 48-well plates pre-coated with 1:20 Matrigel at a density of 40^5^ cells per well in organoid growth medium. After 2 days in culture, monolayers were stimulated with 300ng/ml TL1A (1319-TL, R&D) or 5ng/ml IL-22 (200-22, Peprotech) in a starving medium (Advanced DMEM/F12 with 2mM Glutamax, 10 mM HEPES, 1x N-2 -Gibco-, 1x B-27 without retinoic acid -Gibco-) for 24 hours. Supernatant was collected and RNA of epithelial monolayers was isolated with TRIzol reagent (ThermoFisher).

### Tissue explant cultures

Human intestinal biopsies were washed twice with culture medium (Advanced DMEM/F12, 2mM Glutamax, 10mM Hepes and 200µg/ml Normorcin -InvivoGen-) for 10min on a shaker at RT. Biopsies (one biopsy per condition) were placed in a 48-well plate with 500µl of culture medium with or without 300ng/ml TL1A (1319-TL, R&D). Biopsies were incubated for 20 hours at 37°C. After incubation, biopsies and media were collected on an Eppendorf and spun down for 5 min at 300*g*. Supernatant was further centrifuged for 10min at 16,000*g* and stored at −20°C for further analysis. The tissue pellet was used fresh for RNA isolation.

### RNA isolation

Rectal biopsies preserved with RNAlater or tissue pellets from explant cultures were resuspended in RLT lysis buffer (Qiagen RNeasy Mini Kit) and homogenized in the Bullet Blender 24 (Next Advance) using stainless steel beads (Next Advance) for 7 min at maximum speed. Total RNA was then isolated using the Qiagen RNeasy Mini Kit assay following the manufacturer’s instructions. In contrast, RNA from epithelial monolayers, primary fibroblasts and peripheric CD3+ T cells was isolated with the GeneJET RNA Cleanup and Concentration Micro Kit (ThermoFisher) according to the manufacturer’s instructions. RNA concentration was quantified with the NanoDrop spectrophotometer (Nanodrop technologies), and RNA quality was assessed using the 2100 Bioanalyzer (Agilent). Only samples with RNA integrity number (RIN) > 7.0 were used.

### Bulk RNA-seq experiments

Barcoded RNA-seq libraries were prepared from total RNA of primary intestinal fibroblasts stimulated with TGFβ, LTα_1_β_2_ or TNF (n=6). A TruSeq Stranted mRNA library Perp Kit was used following the manufacturer’s instructions (Illumina, San Diego, CA, USA) at Macrogen (Korea). Libraries were loaded onto a flow cell where fragments were captured on a lawn of surface-bound oligos complementary to the library adapters. Each fragment was then amplified into distinct clonal clusters through bridge amplification. Templates were then sequenced with sequencing by synthesis (SBS) technology.

### Bulk RNA-seq data analysis

From each sample, we performed bulk RNA-seq (paired-end). Each data point was aligned by STAR(16) (v2.7.10a) with GRCh38.p13.genome.fa and gencode.v41.annotation.gtf files. FeatureCounts(17) function was conducted to obtain a count matrix for each gene using Rsubread(18) (v2.16.1) and edgeR(19) (v3.32.1) packages. FPKM was performed for normalization.

The DEG analysis to compare the paired cytokine-treated and control groups was conducted using DESeq2(20) (v1.30.1). The design matrix was constructed as follows: “design = ∼sample_id + treatment”. The signature for each cytokine was obtained by only extracting the upregulated DEG (adjusted p-value < 0.05 and Log2FC > 1.4). We used the genes from TGFβ signature to perform the “AddModuleScore” function as described above in our scRNA-seq data. While comparing the two groups, we performed the Wilcoxon-rank sum test (two-sided) to obtain p values.

The Gene Ontology analysis (biological process) was conducted using “ToppFun” from https://toppgene.cchmc.org/, with the same signature applied to each treatment. FDR < 0.05 (Bonferroni) was used to extract significant pathways.

### External scRNA-seq data analysis of fibroblasts

We obtained scRNA-seq data from CCD-18co human colon fibroblast cell lines treated with various cytokines from GSE233063(21) as CellRanger output. Each CellRanger output file was processed using the Seurat pipeline. We used the same thresholds for mitochondrial gene percentages (< = 25 %) and the number of features (> = 200) as the original paper. After running the Seurat default pipeline, we performed DEG analysis between the control and TGFβ treated samples using the “FindMarkers” function. We extracted a TGFβ signature by using the genes with avgLog2FC > 1.4 and adjusted p-value < 0.05 from TGFβ-treated samples. We used this signature to perform the “AddModuleScore” function as above in our scRNA-seq data. While comparing the two groups, we performed the Wilcoxon-rank sum test (two-sided) to obtain p values.

### Microarray analysis

RNA from IL-22-stimulated 3D epithelial organoids was reverse-transcribed to cDNA using the reverse-transcriptase high-capacity cDNA Archive RT Kit (Applied Biosystems) and hybridized to high-density oligonucleotides using the Affymetrix GeneChip Human Genome U219 Array (Affymetrix, USA). Raw data was then analyzed using Bioconductor tools in R (v3.1.0). We extracted the IL-22 signature by thresholding the adjusted p-value < 0.01. We used this signature to perform the “AddModuleScore” function as described above in our scRNA-seq data. While comparing the two groups, we performed the Wilcoxon-rank sum test (two-sided) to obtain p values.

### Cytosig analysis

We performed the CytoSig analysis to infer which cytokine explains the current gene expression based on the ridge-regression algorithm. We ran the package using “CytoSig_run.py -i [input: scRNA-seq], , -o [output] -r 0 -s 0 -c 1”. As a result, we obtained a Z-score of cytokine activity for each cell. A comparison between the two groups was conducted in the same manner as the “Gene signature analysis”.

### Quantification of gene expression (qRT-PCR)

RNA was reverse-transcribed to cDNA using the reverse-transcriptase high-capacity cDNA Archive RT Kit (Applied Biosystems). qRT-PCR was conducted using the TaqMan Universal PCR Master Mix (Applied Biosystems) in the QuantStudio3 (Applied Biosystems, ThermoFisher). Predesigned TaqMan primers and *ACTB* as endogenous control were used (Supplementary Table 5). Data are expressed as arbitrary units (AU) upon normalization with the CT-value.

### Cytokine measurement by an enzyme-linked immunosorbent assay (ELISA)

Supernatants of stimulated fibroblasts and tissue explants were used for cytokine detection using the DuoSet ELISA Kits from R&D for human MMP1 (DY901B) and IL-22 (DY782), and the human TFPI2 ELISA Kit from MyBioSource (MBS2507217) according to the manufacturer’s instructions.

### Histology analysis

For hematoxylin and eosin (H&E) staining, deparaffinized 3µm sections were stained with Harris hematoxylin, differentiated with saturated lithium carbonate and counterstained with eosin. For immunostaining, sections were submitted to deparaffinization and antigen retrieval with PT Link (Agilent) using Envision Flex Target Retrieval Solution at low pH (Dako). Sections were blocked with 20% horse serum and incubated overnight at 4C with anti-rabbit OLFM4 (14369S, Cell Signaling, concentration 1/100). Signal was revealed with biotinylated anti-rabbit secondary antibody (Vectastain ABC kit), together with the immunoperoxidase chromogen system (Dako). Image acquisition was performed on a Nikon Ti microscope (Japan) using Nis-Elements Basic Research Software (version 5.30.05). Scoring of the staining was done blindly using QuPath (v0.3), where OLFM4+ cells were scored relative to the total number of intestinal crypts.

### Data availability

The raw data from single-cell RNA sequencing is available on GSE290220 and GSE291125. The bulk RNA-seq data from stimulated fibroblasts are available on GSE291014, and the microarray of IL-22 stimulated organoids is available on GSE298981.

### Statistical analysis

Graphs were generated using R (version 4.3.2) and Graphpad Prism (Graphpad Software, version 10). For qRT-PCR analysis of stimulated fibroblasts, epithelial monolayers and tissue explants, fold-change (FC) results relative to the unstimulated condition were log2-transformed and subjected to a One-sample t test. Bulk qRT-PCR analysis of rectal biopsies from the cohort were analysed with Multiple Mann-Whitney t test when comparing CDun vs CD+PFDun patients, while the Kruskal-Wallis test was used when comparing non-inflamed vs inflamed samples from the cohort. A paired t test was used when comparing experimental conditions from the cell line/sample. Two-tailed Pearson correlation was conducted on the log2-transformed FC values of *OLFM4* and *DUOX2* to IL-22 expression.

