## Supplementary Table 1 for "TL1A-activated T cells remodel the rectal mucosa in Crohn’s disease patients with perianal fistulizing disease"

| Study group | CDun | CDinf | CD+PFDun | CD+PFDre | CD+PFDinf |
|---|---|---|---|---|---|
| **Number of patients** | 7 | 5 | 7 | 6 | 6 |
| **Age** | 44 (29 – 68) | 60 (47 - 72) | 54 (29 - 67) | 35 (24 - 59) | 38 (30 – 41) |
| **Sex (male/female)** | 3 / 4 | 2 / 3 | 6 / 1 | 2 / 4 | 3 / 3 |
| **Crohn location** | | | | | |
| • Ileal | 2 | 0 | 4 | 0 | 0 |
| • Ileocolonic | 3 | 1 | 1 | 4 | 5 |
| • Colonic | 2 | 4 | 2 | 2 | 1 |
| ○ Rectum | 0 | 5 | 0 | 6 | 6 |
| **Crohn phenotype** | | | | | |
| • Inflammatory | 2 | 3 | 4 | 3 | 3 |
| • Stricturing | 4 | 1 | 0 | 0 | 0 |
| • Penetrating | 1 | 1 | 3 | 3 | 3 |
| **Treatments** (previous / current) | | | | | |
| - Immunosuppressants* | 4 / 4 | 3 / 0 | 3 / 4 | 5 / 1 | 5 / 1 |
| - Anti-TNF | 3 / 3 | 2 / 0 | 3 / 4 | 5 / 2 | 6 / 1 |
| - Vedolizumab | 1 / 0 | 2 / 1 | 0 / 0 | 0 / 1 | 0 / 0 |
| - Ustekinumab | 0 / 3 | 2 / 0 | 0 / 2 | 1 / 2 | 1 / 3 |
| - Tofacitinib | 0 / 0 | 1 / 1 | 0 / 0 | 0 / 0 | 0 / 1 |
| **Previous perianal surgery** | - | - | 2 | 3 | 3 |
| **Perianal remission** (months) | - | - | 7,2 (6,4 – 12,2) | 17,2 (7 – 20,9) | - |
| **PDAI** | - | - | 2 (0 – 8) | 2 (0 – 9) | 5 (4 – 13) |
| **Partial SES-CD rectum** | 0 | 5 (3 – 9) | 0 | 0 | 5,5 (3 – 7) |
| **CDEIS Rectum** | 0 | 9,35 (7,5 - 36) | 0 | 0 | 16,7 (7 – 24) |
| **Pelvic MRI** | - | - | | | |
| • AGA classification | | | | | |
| ✓ Simple | | | 0 | 2 | 0 |
| ✓ Complex | | | 7 | 4 | 6 |
| • Parks classification | | | | | |
| ✓ Intersphincteric | | | 5 | 5 | 3 |
| ✓ Transphincteric | | | 2 | 0 | 2 |
| ✓ Suprasphincteric | | | 0 | 1 | 1 |
| ✓ Extrasphincteric | | | 0 | 0 | 0 |
| • mVan Assche index | | | 12 (7 – 18) | 6 (2 – 14) | 15,5 (12 – 21) |
| • MAGNIFI index | | | 12 (7 – 18) | 7 (3 – 16) | 15 (13 – 22) |
